# A systematic approach to study protein-substrate specificity enables the identification of Ssh1 substrate range

**DOI:** 10.1101/2022.05.04.490583

**Authors:** Nir Cohen, Naama Aviram, Maya Schuldiner

## Abstract

Many cellular functions are carried out by protein pairs, or families, providing robustness alongside functional diversity. For such processes, it remains a challenge to map the degree of specificity versus promiscuity. Protein-protein interactions (PPIs) can be used to inform on these matters as they highlight cellular locals, regulation and, in cases where proteins affect other proteins – substrate range. However, methods to study transient PPIs systematically are underutilized. In this study we create a novel approach to study stable as well as transient PPIs in yeast. Our approach, Cel-lctiv (<u>CEL</u>lular biotin-<u>L</u>igation for <u>C</u>apturing <u>T</u>ransient *I*nteractions *in <u>V</u>ivo*), uses high- throughput pairwise proximity biotin ligation for uncovering PPIs systematically and *in vivo*. As a proof of concept we study the homologous translocation pores Sec61 and Ssh1. We show how Cel-lctiv can uncover the unique substrate range for each translocon allowing us to pinpoint a specificity determinator driving interaction preference. More generally this demonstrates how CEl-lctiv can provide direct information on substrate specificity even for highly homologous proteins.

## Introduction

A major driver of the evolution of cellular functions is duplication of genes followed by specialization of the homologous pair (Fenech et al., 2020). While gene duplication and specialization broaden functionality, provide cellular robustness and increase regulatory capacity, they pose major challenges when trying to uncover the specialized function of each homologous protein product. The difficulty is increased by the fact that despite exhibiting some specificity, that enables diversification of their functions, they also often demonstrate some degree of functional overlap – thus providing backup for each other (Ihmels et al., 2007). Obviously, this becomes even more challenging as the number of homologs increases and large protein families are formed. Hence it is a general need and goal in cell biology to find approaches that will enable uncovering the unique functional aspects of proteins that have homologs or are part of a protein family. For proteins that act on other proteins, such as kinases, proteases, chaperones and more, these approaches would entail uncovering their exact substrate range as a means to deciphering their specificity or promiscuity.

Methods to study the selectivity of proteins with similar functions often utilize *in vitro* approaches and this requires observing a single (or very few), well behaved, substrates. *In vivo* approaches can also be used and these usually ablate or alter one protein and assay the cellular outcome of this manipulation. However, when one protein is silenced or eliminated, substrates may reroute to back-up pathways (giving rise to false negatives). Alternatively, compensatory rewiring of the cellular system may cause indirect effects (giving rise to false positives). Thus, to unravel substrate range of homologous proteins and reveal the basic principles driving their specialization, there is a clear need for new approaches. These should rely on systematic techniques that would work well for many types of homologs. Such methods should also have the capacity to work *in vivo* and provide information on cellular activity preference without deleting the tested protein itself.

One powerful way to uncover the specialized cellular functions that differentiate homologous proteins is by assaying their stable, and transient, interactions. Knowing these interactions may suggest different cellular contexts in which these proteins work (For example different cellular locals or different complex members), unique regulatory mechanisms (such as various post translational modification enzymes working on them) and, for the subgroup of proteins working on other proteins as substrates, also their unique and specific substrate range.

In this study we utilize the yeast, *Saccharomyces cerevisiae* (from hereon called simply yeast) to create a novel, systematic, approach to measure the differential stable and transient interactomes for proteins that govern parallel functions as a means to map their functional specificity. Our approach uses high- throughput pairwise proximity biotin ligation assays for uncovering protein-protein interactions systematically and *in vivo*. We have named it Cel-lctiv (pronounced like “selective” - <u>CEL</u>lular biotin-<u>L</u>igation for <u>C</u>apturing <u>T</u>ransient *I*nteractions *in <u>V</u>ivo*). Cel-lctiv overcomes the limitation of hypothesis driven, small number of candidates by assaying the entire proteome and the false positive/negative issues around loss of function by maintaining the complete cellular context.

As a proof of concept, we chose to use the two homologous translocation pores into the endoplasmic reticulum (ER) – together catering for over 30% of the proteome that must be translocated into the secretory pathway. The well-studied translocation pore is the Sec61 translocon (Bieker & Silhavy, 1990; Deshaies et al., 1991), while the less studied homolog is Ssh1 (Sec Sixty-one Homolog 1) (Finke et al., 1996). Since translocation through a pore is an extremely transient event and since Sec61 and Ssh1 have overlapping functions they have therefore provided a challenge for uncovering the repertoire of translocating substrates in the past. Since the presence of a Sec61 homolog is conserved from yeast to humans (The human SEC61A1 homolog is termed SEC61A2), it stands to reason that there is an evolutionary purpose or advantage for maintaining the pair however this has not yet been uncovered.

In the past, attempts to uncover the reason for having two homologous translocons have focused on hypothesis driven, single candidates and measured their translocation capacity either *in vivo* or *in vitro*. These classic studies have been extremely important in providing the first candidate substrates for Ssh1 (Aviram et al., 2016; Spiller & Stirling, 2011; Wittke et al., 2002). However, from this handful of potential candidates, each having distinct properties and no clear unifying aspect, it has been difficult to determine the rules governing specificity. Beyond the small number of potential proteins tested to date, an additional complication has been the use of loss of function approaches. Since Sec61 is an essential protein but Ssh1 is not (albeit its loss renders the cells extremely slow growing (Wilkinson et al., 2001)), most such experiments were done uni- directionally on cells deleted for Ssh1. This has led to the misconception that Ssh1 is simply a backup translocon with potentially only a small number of unique substrates none of which are essential under normal growth conditions.

By combining Cel-lctiv to assay the interaction preference of both Ssh1 and Sec61 coupled with whole proteome analysis on the effects of *ssh1* loss, we uncover the range of unique cellular effects of losing Ssh1 as well the degree of functional overlap between Sec61 and Ssh1 and their unique potential substrate range. Moreover, having a wide range of new substrates unique for each translocon allowed us to decipher a biochemical specificity determinator for their interaction preference. More generally this demonstrates how Cel-lctiv can provide direct information on substrate selectivity and specificity even for highly homologous proteins and does this in the native cellular environment on minimally perturbed proteins and in a systematic fashion.

## Results

### Development of Cel-lctiv - Cellular biotin-Ligation for Capturing Transient Interactions in vivo

To uncover the different cellular environment, regulatory pathways and substrates of a protein machinery it is essential to capture not only their stable interactions but also the very transient ones. For example, when trying to understand the distinct roles of the two translocons Sec61 and Ssh1 it would be important to understand which proteins are transiently translocating through them in an *in vivo* setting (Figure 1A). While assaying transient interactions between two proteins is feasible (Villalobos et al., 2007), it has remained difficult to do this systematically on a proteome wide level. Hence, we developed a new approach, that we call Cel-lctiv (Cellular biotin-Ligation for Capturing Transient Interactions *in vivo*).

**Figure 1:**
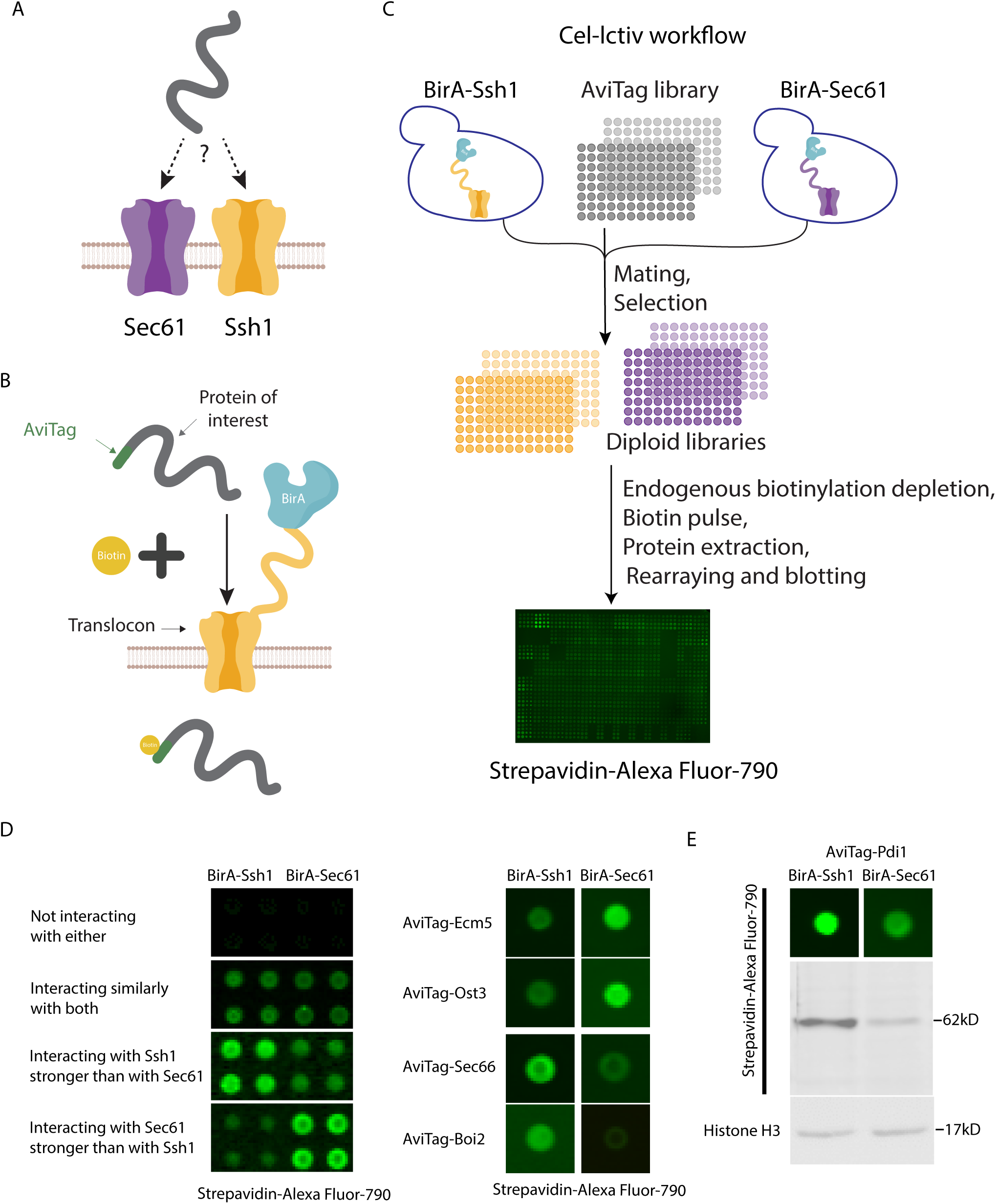
Cel-lctiv - Cellular biotin-Ligation for Capturing Transient Interactions in vivo – is a powerful approach for measuring protein specificity. A) A schematic illustration of the research question – how can we define differential substrate range for two homologous proteins? Shown as an example are the two endoplasmic reticulum translocons Sec61 and Ssh1. B) A schematic illustration of the Cel-lctiv approach to uncover the repertoire of stable and transient protein protein interactions. When a protein (in this case one of the translocons) is tagged with the specific biotin ligase BirA and the potential substrate is tagged with the biotin acceptor AviTag even a transient physical interaction will allow biotinylation of the AviTag and this can be readout using fluorescent streptavidin. C) A schematic illustration of the work flow of Cel-lctiv: Strains containing the BirA tagged proteins (in this case either one of the two translocons) are mated with a collection of ∼6000 yeast strains in each of which one protein is N terminally tagged whit AviTag under the native promoter and localization signal. Each resulting diploid strain undergoes automated protein extraction, blotting onto a membrane and visualization using fluorescent streptavidin. Biotinylation levels are compared between the two samples to identify an interaction preference. D) Cel-lctiv results were clustered into groups: proteins that did not show a detectable interaction with either translocon, proteins that showed similar interaction with both translocons, and proteins that showed a preference for one of the translocons. Examples of the latter groups are shown. E) Comparison between the dot-blot and Western blot analysis of the same lysate from the screen to verify that the signal difference results from differential substrate biotinylation.

Cel-lctiv relies on proximity labeling of a biotin ligase, BirA, to its specific acceptor peptide AviTag (Beckett et al., 1999). However, instead of doing this for a single pair of proteins, Cel-lctiv measures the pair-wise interaction abundance for a given protein with every other protein in the yeast proteome. The benefit of using such an approach is that each interaction event results in a separately measured output so the sum of interactions can be quantified to provide a global analysis of biotinylation levels per protein pair. Comparing the same AviTagged substrate with different BirA tagged proteins (for example two homologs) allows the direct comparison of their relative interaction preference (Figure 1B). Unlike other non-specific biotin ligation methods, such as TurboID, there is only a single, identical, ligation site for each protein enabling a linear relationship between the number of molecules that have interacted and the signal intensity. In addition, since this is performed in a pair-wise manner it allows the detection of low abundance proteins that could be masked in pooled approaches. This is especially important when trying to uncover a complete substrate range.

One major difficulty with using the pair-wise biotinylation as a robust tool, especially if relying on endogenous expression levels as we have done (see below), is the high background of endogenous biotinylation in yeast. To reduce the background biotinylation we have therefore utilized our previously established methodology for reducing endogenous biotinylation, ABOLISH (Auxin- induced BiOtin LIgase diminishing) (Fenech et al., 2022). This allowed us to reduce background noise and increase our sensitivity especially for low-abundance proteins.

To develop Cel-lctiv we utilized two recently established yeast strain collections (libraries). In one library each protein in the yeast proteome is tagged with an N terminal AviTag and expressed under its endogenous promoter and localization signal (such as signal peptide (SP) or mitochondrial targeting signal (MTS)) if it exists. In the second library each protein is similarly tagged with an N terminal BirA retaining the natural promoter and targeting information (Fenech et al., 2022). To perform Cel-lctiv you first choose your protein pair or family from the BirA library (we selected BirA-Sec61/Ssh1). Then the BirA strains are mated with the whole-genome Avitag collection that is of the opposite mating type. This generates new diploid whole-genome collections carrying both selected traits. In our case each contains one of the translocons (Ssh1 or Sec61) tagged with BirA and all yeast proteins tagged with AviTag (Figure 1C).

To enable the quantitation of thousands of pair-wise interactions in quadruplicate we developed a high-throughput dot blot approach using fluorescent streptavidin to visualize the degree of biotinylation in each strain following the background biotinylation reduction by ABOLISH (Figure 1C). The result is then computationally analysed to measure relative biotinylation signal for each BirA and AviTag combination.

Focusing on our protein pair, Sec61 and Ssh1, we found that those that did not interact with either, that interacted with both and, using a cut-off of a two-fold higher signal, those that favour a specific translocon over the other. Specific substrates include Ecm5 and Ost3 interacting preferentially with Sec61 while Sec66 and Boi2 interacted preferentially with Ssh1 (Figure 1D). To quality control our method we remeasured a select set of protein extracts from the original assay by resolving them on a SDS-PAGE gel thus allowing us to assay the specific AviTag biotinylation contribution to the total biotinylation and ensuring proper molecular weight as expected from that strain (Figure 1E).

Collating the complete set of proteins (Supplementary Table 1) we found that of the previously suggested substrate of Ssh1, the ones that we could assay, Sec66 (Spiller & Stirling, 2011) and Prc1 (Wilkinson et al., 2001) both indeed showed a preference for Ssh1 supporting the ability of Cel-lctiv to uncover the transient interactome of two homologous proteins.

### Cel-lctive provides a global insight into Ssh1 substrate selectivity

Looking globally at the whole proteome we found that, as expected, the majority of proteins did not interact with either translocon (2742, note that in this group we cannot differentiate between biological and technical reasons). From the interacting proteins many had similar signal levels for both translocons suggesting a high degree of functional overlap (1674, out of which 428 are potential substrates). Importantly, our Cel-lctiv approach uncovered many proteins that have a preference for one translocon. Out of those we reasoned that direct substrates would be those that have a SP or a signal anchor (a transmembrane domain (TMD)) – either one of which is required to engage the pores for productive translocation. The screen uncovered 111 proteins that showed a preference for Ssh1, 30 of which contain a SP or at least one TMD (from these we removed those that have an MTS, since those could be mitochondrial inner membrane proteins) - making them potential secretory substrates. 285 proteins showed a preference for Sec61, 72 of them potential secretory substrates (Figure 2A, Supplementary Table S1). This already suggests that while Ssh1 and Sec61 can indeed act as back-ups for each other for the vast majority of potential substrate proteins, they also each have a unique function since tens of proteins preferred to translocate specifically through one or the other under the ploidy and growth condition that we measured. Amidst the biotinylated proteins we also found differential interactions with potential regulators such as the kinases Gin4, Pkc1, Ptk2 and Kdx1 interacting with Ssh1 in comparison to Bck1 and Iks1 interacting with Sec61. These differential kinase- translocon interactions suggest that they are either affected by or affect different regulatory networks (Supplementary Table S1). Surprisingly, 12.6% of the interacting proteins of Ssh1, were from organelles that are thought to be separate from the secretory system, such as mitochondria (Supplementary Table S1).

**Figure 2:**
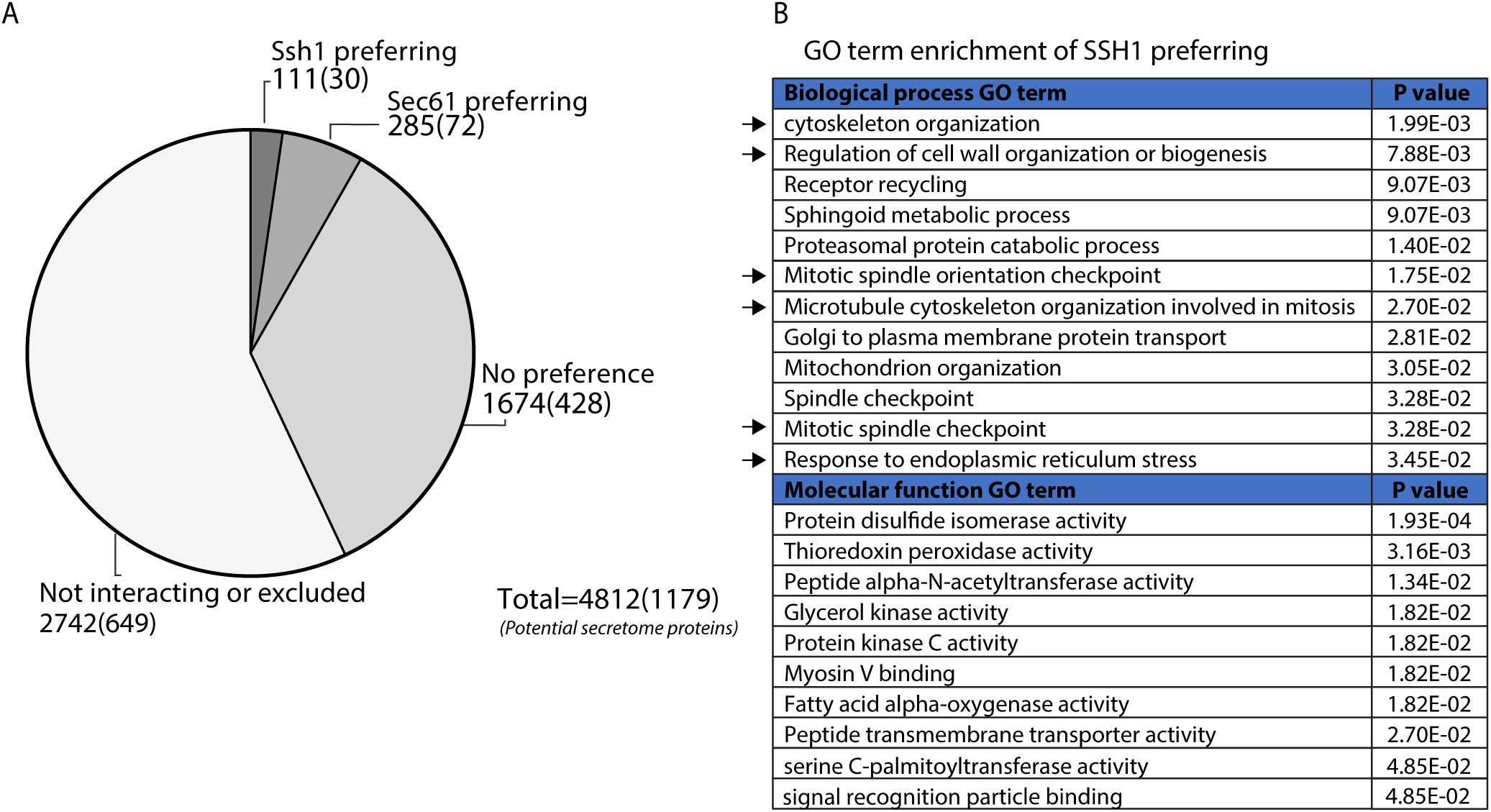
Cel-lctive provides a global insight into Ssh1 substrate selectivity. A) A pie chart showing the distribution of Cel-lctiv results divided into non interactors/excluded, equally interacting with both or differential preference proteins. Numbers represent all proteins that fit into the group with those in parentheses referring to the subgroup that could be direct substrates (predicted to have a SP or at least one TMD and no MTS). B) GO term enrichment analysis of proteins that showed an interaction preference with Ssh1, using Yeastmine. Black arrows indicate terms that intersect with the GO term enrichment analysis of *Δssh1* affected proteins (Figure 4).

To assay for cellular processes that are overrepresented in the group preferentially interacting with Ssh1, we turned to Gene Ontology (GO) term enrichment analysis (Cherry et al., 2012; Engel et al., 2014) (Figure 2B, Supplementary Table S2). The analysis was enriched with the expected GO terms that describe the core translocation machinery such as “Signal recognition particle binding” and “peptide transmembrane transporter activity”. However, some unexpected GO terms also came up as enriched highlighting processes related to cytoskeleton organization, cell wall biogenesis and response to ER stress (Figure 2B). Altogether these suggest that there is a functional divergence of the specific substrates that utilize Ssh1.

### A high-throughput screen uncovers the global effect of Δssh1 on the proteome

The Cel-lctiv approach enabled us to find potential translocating substrates that prefer one translocon over another. However, this gives little insight as to what the physiological outcome of this preference is. One way to assess the physiological effects of having non overlapping functions for the translocons is to gauge the cellular effects of losing one of them. To uncover the global cellular effects, those that cannot be buffered by the presence of a homolog, we focused on the non-essential translocon Ssh1. To do so we visualized all yeast proteins (tagged on their N terminus with a Green Fluorescent Protein (GFP) under the *NOP1* promoter (Weill et al., 2018; Yofe et al., 2016), on the background of either a *Δssh1* or a control using a high-throughput confocal spinning disc imaging platform (Figure 3A). We manually analyzed the images and categorized proteins as affected if the protein changed signal localization and/or shape in the mutant relative to the control. We found 252 proteins that changed their localization (as exemplified by GFP-Yck1 (Figure 3B)), 170 proteins that displayed a specific type of partial change of localization with the appearance of punctate structures (as exemplified by GFP-Yet3 (Figure 3B)) and 65 proteins that displayed a complete change in their localization (as exemplified by GFP-Ndj1 (Figure 3B)) (for an overview of all changes see Figure 3C and Supplementary Table S1). Altogether over 10% of the assayed proteome was affected by Ssh1 loss suggesting a significant and global effect of its loss as well as rewiring of the cell to accommodate loss of this translocation channel. This highlights its unique cellular role and its non-redundant functions.

**Figure 3:**
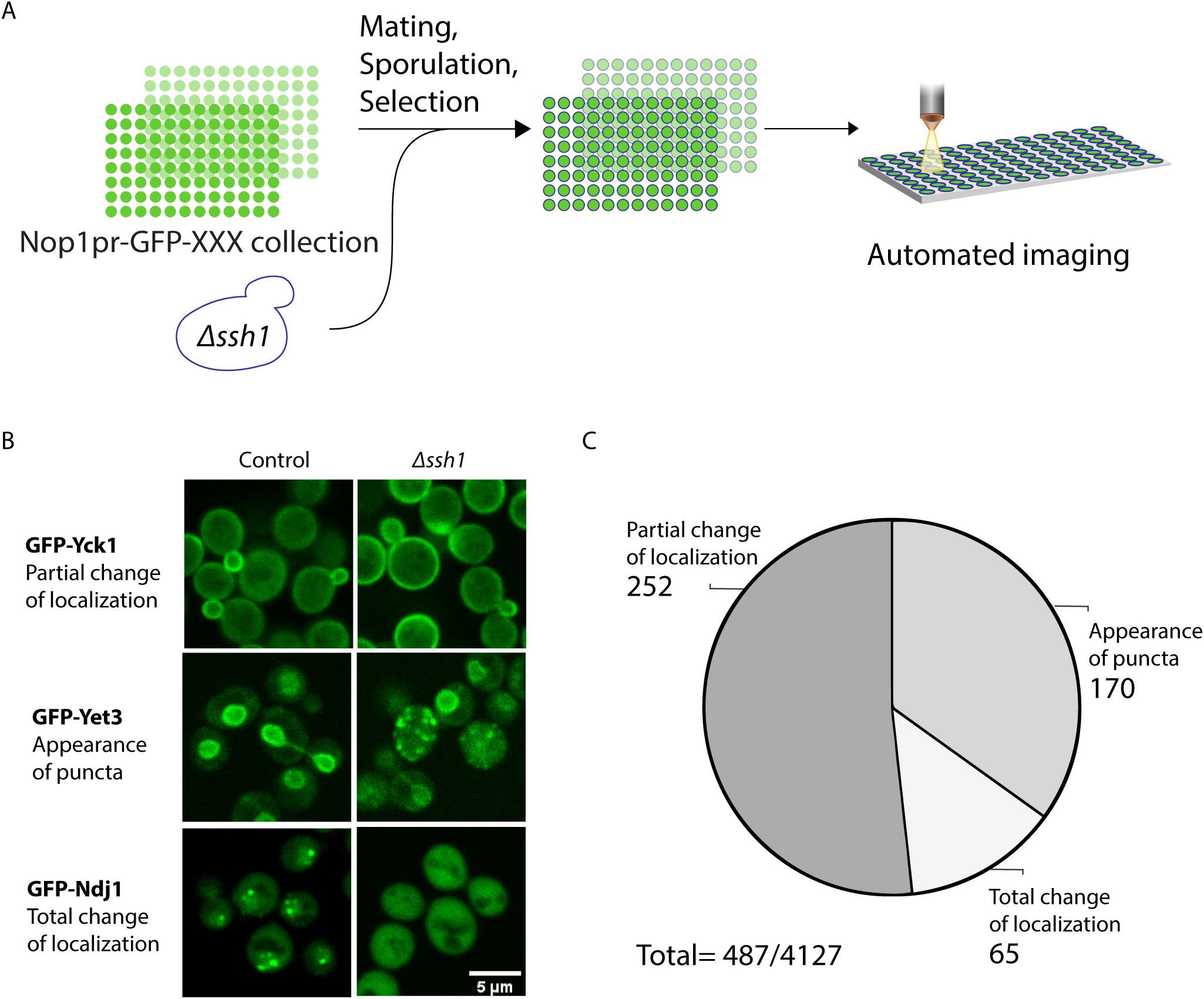
A high-throughput screen uncovers the effect of Δssh1 on protein localization. A) A schematic illustration of the high-throughput screen aimed to identify the cellular effects of losing Ssh1. A strain where *ssh1* was deleted was integrated into a library of ∼6000 yeast strains in each of which one protein is N terminally tagged with GFP under the constitutive *NOP1* promotor. Strains were automatically imaged using a high-throughput spinning-disk confocal microscope and manually examined to uncover proteins that show a change in signal shape and/or localization. B) Representative strains from the three groups into which all hits were divided: Proteins demonstrating a partial change in localization, proteins that showed an appearance of puncta, and proteins that showed a total change in localization. Scale bar=5µm. C) Pie chart showing the overall results from the high-throughput screen divided into the three categories from B (for the complete list of hits see Supplementary Table S1).

### Transient interactors that showed a preference for Ssh1 suggest the mechanistic link to its cellular effects

While the above results show global changes, we wished to uncover the major cellular processes that were affected by loss of Ssh1. To assay for cellular processes that are overrepresented in the proteins whose localization was altered by loss of Ssh1, we turned again to GO term enrichment analysis (Cherry et al., 2012; Engel et al., 2014). The analysis showed enrichment for a variety of processes related to transport and cytoskeleton organization. However, the most consistent and repeating GO term group was those related to budding and polarity process. These include: “Establishment or maintenance of cell polarity”; “Development process involved in reproduction”; “Bipolar cellular bud site selection”; “Cell budding” and “Structural constituent of cell wall” (Figure 4A, Supplementary Table S2).

**Figure 4:**
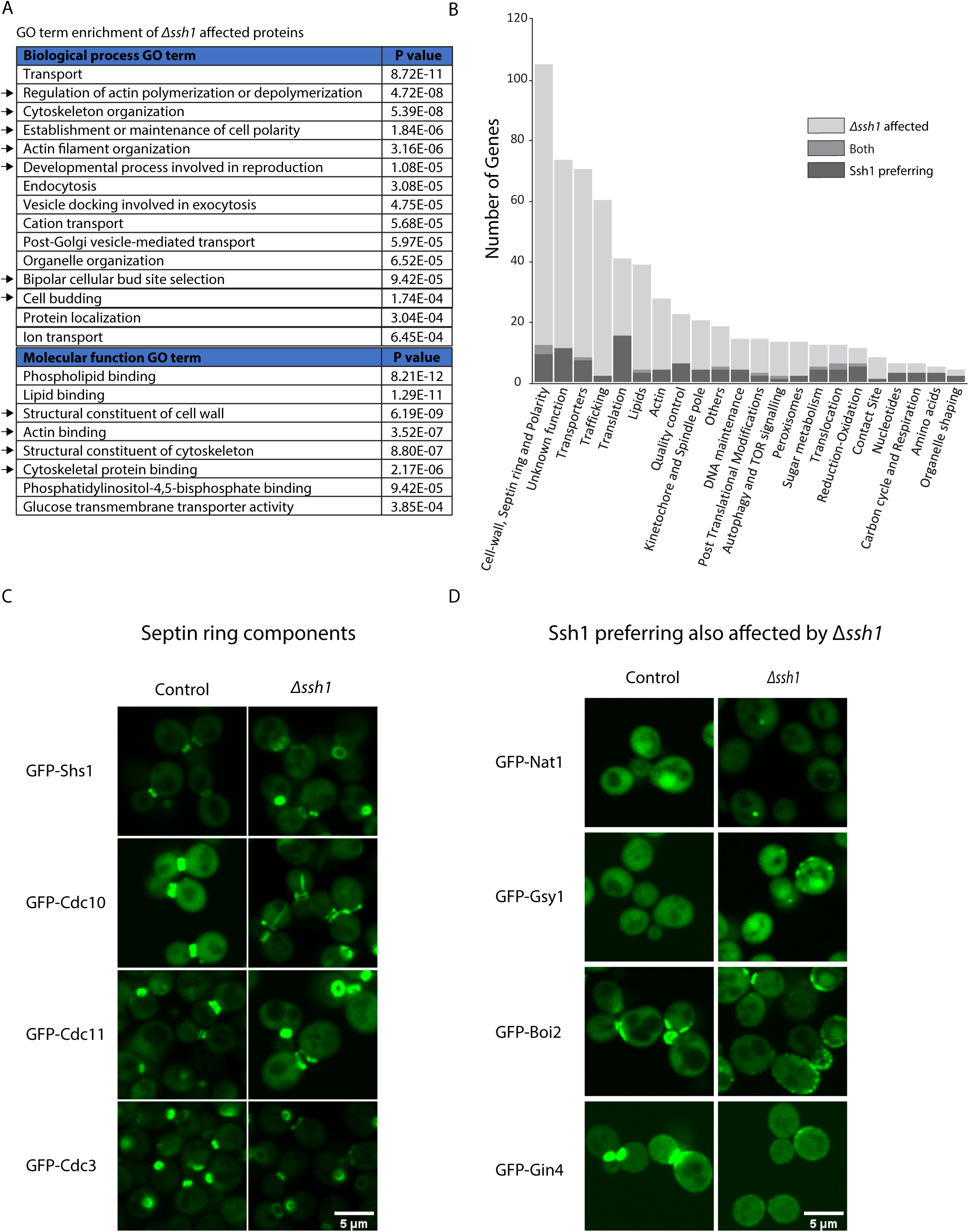
Loss of Ssh1 leads to defects in Septin ring morphology. A) GO term enrichment analysis of all proteins affect by the *Δssh1* background. Black arrows indicate terms that intersect with the GO term enrichment analysis of proteins that showed an interaction preference with Ssh1 (Figure 3). B) A bar graph of the manually assigned functional groups showing crosstalk between the two screens. C) Microscopy images of N terminal GFP tagged Septin ring components in the *Δssh1* and control backgrounds. The comparison shows accumulation of open ring forms and alterations in Septin ring structure in the mutant. Scale bar=5µm. D) An example of proteins that came up in both screens hence both showed an interaction preference and a localization effect in the *Δssh1* background. Scale bar=5µm.

Since GO terms are limited in their functional information, and to be able to compare the outputs of the two systematic screens (uncovering either proteins that preferentially bind Ssh1 and those that are affected by its loss), we manually grouped the protein hits from the two screens into functional categories based on the description and annotation in the yeast genetics database (SGD) (Cherry et al., 2012; Engel et al., 2014) (Supplementary Table S1). These were then used to group the hits from the two approaches and visualize those aspects of cellular function that were most represented from both approaches together. Our analysis uncovered that the most enriched groups were “Cell wall, Septin ring and polarity”; “Transporters”; “Trafficking” and “Translation”. The largest group of proteins represented in both screens were in the “Cell wall, Septin ring and polarity” and, as expected, in “Translocation” (Figure 4B, Supplementary Table S1).

The major structure that represents the outcome of many of the above cellular processes (cytoskeleton, polarity and budding) is the Septin ring. The Septin ring is a protein scaffold that assembles on the dividing septum (bud neck) during cell division and is involved in the selection of the bud-site, the positioning of the mitotic spindle, polarized growth, and cytokinesis (Longtine & Bi, 2003; Oh & Bi, 2011).The Septin structure matures throughout the cell cycle: it first appears as a distinct ring; after bud emergence the ring broadens to assume the shape of an hourglass around the mother-bud neck; finally during cytokinesis the Septin cortex splits into a double ring which eventually disappears. The clear visual structure of the Septin ring prompted us to follow the major Septin ring components in *Δssh1* strains. Indeed, we found that for all major Septin subunits that we assayed (Shs1, Cdc10, Cdc11 and Cdc3) there was a clear enrichment for the open, ring, stage and the ring was also enlarged relative to control cells (Figure 4C). This may be an indication of delayed cytokinesis which would also explain the reduced growth rate of *Δssh1* strains (Wilkinson et al., 2001). Such a delay may be due to the activation of the ER surveillance stress response (ERSU) (Piña & Niwa, 2015) as result of the UPR activation in *Δssh1* (Wilkinson et al., 2001). The ERSU monitors the health of the ER and blocks cytokinesis in cells that have stressed ER as a way to prevent its inheritance.

Indeed, of the 12 proteins that came up in both screens were Boi2 and Gin4, that are directly related to the budding process. Gsy1, a glycogen synthase whose expression was shown to be induced by osmotic shock (Unnikrishnan et al., 2003), and the Nat1 N-acetyltransferase. Together they suggest a mechanistic link between direct Ssh1 interaction and effects on cell wall. This suggested that specific direct interactors of Ssh1 can explain the physiological effects of its absence on Septin ring formation.

### The biochemical properties of the signal peptide determine translocon preference

If proteins exist that have a translocation preference then there should be sequence determinants that encourage translocon choice. Having a list of potential substrates that prefer interacting with (and therefore potentially translocating through) one translocon or another also allowed us to explore the biochemical properties that differentiate Ssh1 and Sec61 substrates. We focused on those interactors that contains a SP due to its role in translocon engagement. For these substrates we explored the biochemical properties of the SP by comparing the hydrophobicity (using a Kyte- Doolitle scale) across the various positions in the first few amino acids of proteins that prefer one translocon, the other or have no clear interaction preference and found that Ssh1 interactors have distinct hydrophobicity in their most N terminal residues (Figure 5A).

**Figure 5:**
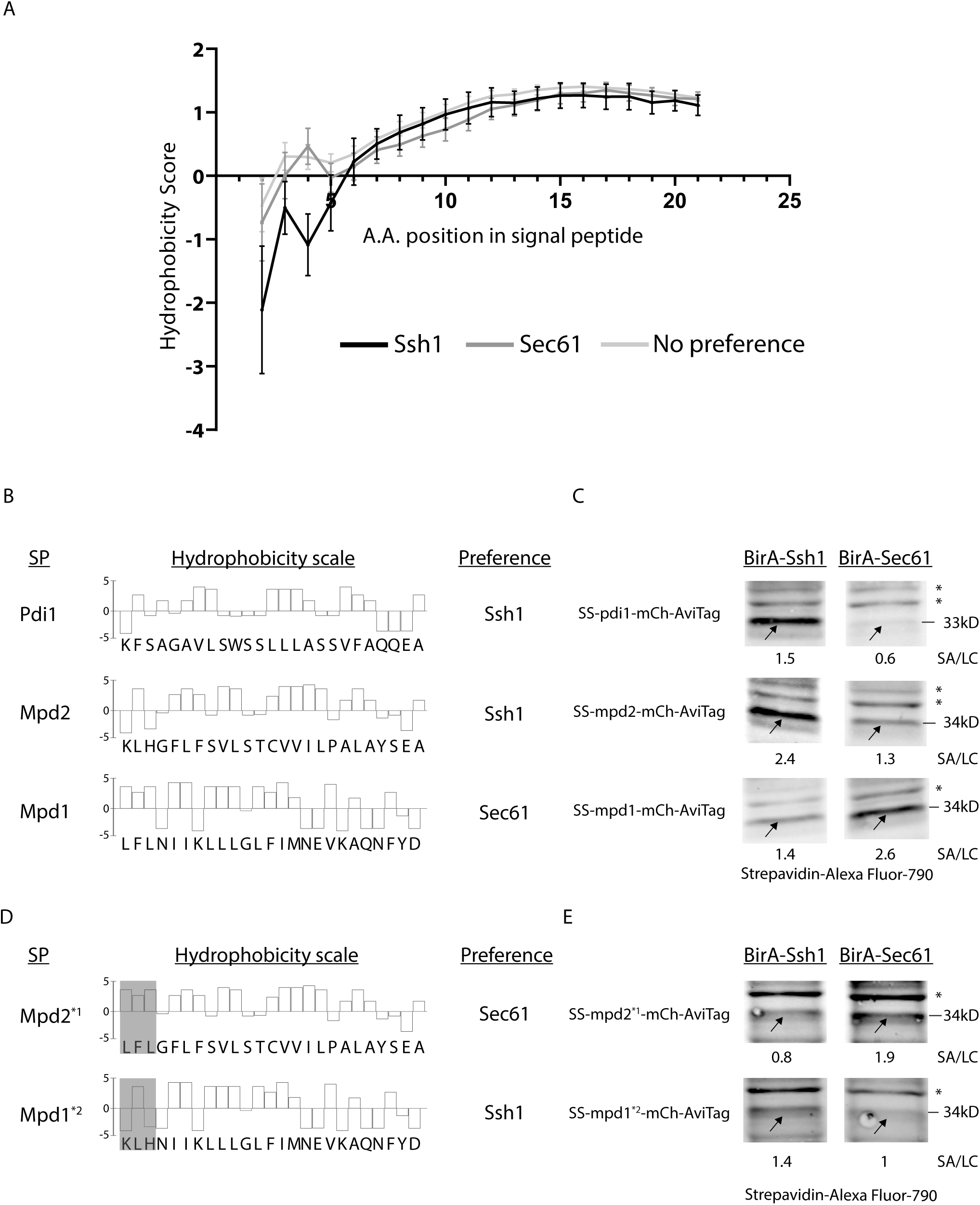
The biochemical properties of the signal peptide determine translocon preference. A) A graph showing the Kyte-Doolittle hydrophobicity scale average for all predicted SPs from the group of proteins that showed an interaction preference to either Ssh1, Sec61, or that interacted with both similarly. B) A bar graph showing the Kyte-Doolittle hydrophobicity scale for the predicted SPs of PDI family members that came up as differential interactors in our Cel-lctiv assay. Shown are Pdi1 and Mpd2, two PDI family members that showed an interaction preference to Ssh1, Mpd1 that showed an interaction preference to Sec61, C) Western blots showing the biotinylation level of the SP reporter construct on a Ssh1-BirA or Sec61-BirA background. These highlight that the SP is sufficient to endow interaction preference in the absence of the complete protein context. The SA/LC ratio represent the biotinylation signal (band marked by an arow) and the H3 loading control band total signal ratios. Asterisks (*) mark non-specific bands. D) A bar graph showing the Kyte-Doolittle hydrophobicity scale for the two variants of Mpd1 and Mpd2 where the three first amino acids were swapped aiming to alter the interaction preference. E) Western blots showing the biotinylation level of the two variants of Mpd1 and Mpd2 SP reporter construct on a Ssh1-BirA or Sec61-BirA background. Showing that swapping the first three amino acids is sufficient to convert the SP reporter construct interaction preference suggesting that the information for translocon choice is found at the very N terminus of the proteins. The SA/LC ratio represent the biotinylation signal (band marked by an arow) and the H3 loading control band total signal ratios. Asterisks (*) mark non-specific bands.

To look at this in more detail we chose to study three proteins that belong to the Protein Disulfide Isomerases (PDI) family. The PDI family contains five proteins (Nørgaard et al., 2001) three of which we could detect in our assay. Surprisingly all three, Pdi1, Mpd2, and Mpd1, showed an interaction preference for one of the translocons. Pdi1 and Mpd2 showed preference for Ssh1 while Mpd1 showed a preference to Sec61 (Supplementary Table S1). Indeed, looking into their SPs they each had hydrophobicity patterns that suited the preference shown by the group analysis (Figure 5B). Two PDI family members (Eps1 and Eug1) were not represented in the screen at all, however their SP pattern suggests that they too are Ssh1 substrates.

To assay whether the observed SP properties are sufficient to drive the interaction preference, we generated a construct that contains only the SP of either Pdi1, Mpd1 and Mpd2, fused to a reporter containing an mCherry and an AviTag. By assaying the interaction preference on the background of either Ssh1-BirA or Sec61-BirA we could observe that indeed the SP alone was sufficient to generate an interaction preference similar to that observed by the full protein (Figure 5C). Hence small differences in SP biochemical properties underly translocon selectivity.

To specifically demonstrate that the information that lies in the SP is found in the first amino acids we performed a domain swap experiment in which we exchanged the first three amino acids of the Mpd1 and Mpd2 SP (creating the constructs Mpd1^*2^ and Mpd2^*1^) (Figure 5D). Indeed, simply exchanging these amino acids was sufficient to alter the translocon preference of a protein (Figure 5E).

Our approach shows how Cel-lctiv can provide a global view of transient interactions on a global scale. It also demonstrates how having the broad knowledge of potential substrate range enables the mining for the specific sequence determinants that govern selectivity. Moreover, while the variability of the SP was previously shown to affect targeting pathway choice and the capacity to open the translocon channel (Johnson et al., 2013) this is the first demonstration that information in the SP also dictates the choice of translocons and by that, potentially the downstream fate of secretory proteins by different sets of accessory proteins, modifications and trafficking.

## Discussion

Uncovering the specificity of protein homologs is a challenging task (Herzig et al., 2012; Megyeri et al., 2019; Yifrach et al., 2016). In this manuscript we have brought forward a new approach for exploring the substrate range for molecular machineries that have proteins as their substrates. Our approach, Cel-lctiv, relies on proximity ligation in a pair-wise setting that enables systematic identification of transient interactions that would be expected from a protein substrate.

We focused on an enigmatic pair – the two homologous SEC translocons into the ER – Sec61 and Ssh1. Using both systematic loss of function studies to gauge the effect of Ssh1 loss on all cellular proteins as well as discerning the differential interactors of Sec61 and Ssh1 we could not only uncover the repertoire of cellular processes affected specifically by Ssh1 loss but also validate the biochemical properties of the SP as one determinant enabling differential substrate selection.

Why would some proteins evolve to use Ssh1 and others Sec61? An intriguing concept that came out of our analysis is that large protein families may have evolved to distribute their entry between the two translocons to increase robustness of their function. One such example is the PDI family of proteins – out of 5 members of this family the three that we could measure had distinct preference for either one of the two translocons. Loss of targeting for two of the PDI family members that prefer Ssh1 may explain why its loss leads to activation of the ER Unfolded Stress Response (UPR) (Wilkinson et al., 2001) and the resulting higher sensitivity to the reducing agent Dithiothreitol (DTT) (Rand & Grant, 2006) as well as to the glycosylation inhibitor tunicamycin (Kapitzky et al., 2010).

An additional hypothesis as to why cells have evolved two entry routes into the ER is that Ssh1 and Sec61 reside in different subdomains of the ER membrane endowing their substrates with a unique post translational environment in which to mature. Another reason may be the capacity to regulate differential entry upon changes in cellular conditions - this is hinted upon by the differential set of regulatory proteins that each translocon interacts with. However, to directly test this hypothesis, our Cel-lctiv approach would have to be performed under additional metabolic or stress conditions. Either way, it is clear that Ssh1 is not simply a back-up translocon and has evolved a unique substrate range and regulatory network.

Surprisingly, our analysis uncovers a huge enrichment for cell wall biogenesis and integrity components as both being affected by the loss of *ssh1* and directly interacting with Ssh1. The direct interactions suggest that maybe specific cell wall proteins prefer to utilize Ssh1 as their primary translocon. This includes the Chitin Synthase Csh2 and the putative chitinase Cts2. Another aspect is the potential regulatory role of Ssh1 by binding, anchoring or modulating regulatory factors such as the bud growth and polarity inducing kinases Gin4, Kdx1 and Pkc1 or phosphatase Msg5 and the Boi1/2 polarity factors required for polarized vesicle fusion.

Another unexpected finding is the presence of mitochondrial membrane proteins as specific interactors of Ssh1. One such protein, Tim21 is a mitochondrial inner membrane protein that both interacts with Ssh1 and is affected by its loss. Tim21 is a typical substrate of the ER-SURF pathway in which mitochondrial inner membrane proteins utilize the ER surface to target to mitochondria (Hansen et al., 2018; Koch et al., 2021). Indeed, strains deleted for *SSH1* have previously been shown to have respiratory effects, enhanced DNA loss (Wilkinson et al., 2001) as well as altered mitochondrial biogenesis (Laborenz et al., 2019).

More generally, our work shows that Cel-lctiv can be used for the study of protein-protein interactions on a whole proteome level and is especially suited for uncovering short, transient interactions in a native environment. This method, available to all (all libraries and protocols are distributed freely) and not requiring sophisticated machinery such as mass spectrometers, is especially powerful for uncovering the substrate range of proteins that work on other proteins. In this respect Cel-lctiv can enable the dissection of substrate range and specificity of protein pairs, protein families or even just groups of proteins undertaking parallel functions. These include kinases/ phosphatases; other post-translational modification enzymes (ubiquitin ligases, glycosyl transferases etc…); proteases and many more. By uncovering the entire repertoire of differentially interacting substrates it should be possible to start dissecting the rules governing specificity and promiscuity of proteins, previously difficult to study.

## Materials and Methods

### Yeast strains and plasmids

*S. cerevisiae* strains were based on the laboratory strain BY4741 (Baker Brachmann et al., 1998). Genetic manipulations were performed using the lithium acetate, polyethylene glycol, single-stranded DNA method (Gietz & Woods, 2006). Plasmids for PCR-mediated homologous recombination were previously described (Janke et al., 2004; Longtine et al., 1998). Supplementary Table S3 lists the primers, plasmids and strains used in this study.

### Western blots

#### Culturing and protein extraction

5ml of cells at 0.5OD_600_ were collected by centrifugation at 3,000g for 3min, washed with 1ml of Double Distilled Water (DDW), resuspended in 200µl lysis buffer containing 8M urea, 50mM Tris, pH 7.5, and oComplete Protease Inhibitors (Merck) and lysed by high-speed bead beater with glass beads (Scientific Industries) at 4 °C for 10min. 25µl of 20% Sodium Dodecyl Sulfate (SDS) was added before incubation at 45°C for 15min. The bottom of the microcentrifuge tubes was then pierced, loaded into 5ml tubes, and centrifuged at 4,000g for 10min to separate the lysate from the glass beads. The flow-through collected in the 5ml tubes was transferred to a fresh 1.5ml microcentrifuge tube and centrifuged at 20,000g for 5min. The supernatant was collected, and 4x SDS-free sample buffer (0.25M Tris, pH 6.8, 15% glycerol, and 16% Orange G containing 100mM DTT) was added to the lysates, which were incubated at 45°C for 15min.

#### Resolving, blotting and acquisition

Protein samples were separated by SDS-PAGE using a 4–20% gradient gel (Bio-Rad) and then transferred onto a 0.45µm nitrocellulose membrane (Pall Corporation) using a Trans-Blot Turbo transfer system (Bio-Rad). Membranes were blocked in 2% wt/vol Bovine Serum Albumin (BSA) in Phosphate Buffered Saline (PBS) solution for 30min at Room temperature (RT), incubated for 1hour at RT with rabbit anti-Histone H3 (ab1791, 1:5,000; Abcam) diluted in a 2% wt/vol BSA/PBS solution containing 0.01% NaN3. After washing, membranes were then probed with secondary goat anti-rabbit -IRDye680RD antibody (ab216777; Abcam) and streptavidin- Alexa Fluor790 (S11378; Invitrogen), both diluted 1:10,000 in 2% wt/vol BSA/PBS solution for one hour at RT. Blots were washed and imaged on the LI-COR Odyssey Infrared Scanner. Images were quantified using GelAnalyzer 19.1 (Lazar & Lazar) high-throughput microscopy screening

##### Library preparation

The synthetic genetic array (SGA) method was used for integrating the desired genomic manipulations into yeast libraries (Cohen & Schuldiner, 2011; Tong & Boone, 2006). Query strains for screens were constructed on a SGA ready strain (YMS721; (Breslow et al., 2008)), and libraries were handled using a RoToR bench-top colony arrayer (Singer Instruments). Briefly, query strains were mated with strains from the library on rich medium plates to generate diploid cells. Cells were then transferred to nitrogen starvation media for one week to induce sporulation. Haploid cells were selected using canavanine and Thialysine (Sigma-Aldrich) lacking leucine to select for MATalpha. The final library was generated by selecting for the combination of manipulations desired. Representative strains from the final library were validated by both microscopy and check-PCR.

##### Culturing and microscopy

Cells were moved from agar plates into liquid 384-well plates using the RoToR bench-top colony arrayer (Singer Instruments). Liquid cultures were grown overnight in synthetic medium with 2% glucose (SD) in a shaking incubator (LiCONiC Instruments) at 30°C. A Tecan freedom EVO liquid handler (Tecan), which is connected to the incubator, was used to back-dilute the strains to ∼0.25OD_600_ in plates containing the same medium. Plates were then transferred back to the incubator and were allowed to grow for 4hour at 30°C to reach logarithmic growth phase. The liquid handler was then used to transfer strains into glass-bottom 384-well microscope plates (Brooks Bioscience) coated with Concanavalin A (Sigma-Aldrich) to allow cell adhesion. Wells were washed twice in a low fluorescence synthetic medium (Formedium) to remove floating cells and reach a cell monolayer. Plates were then transferred into the automated microscopy system using a KiNEDx robotic arm (Peak Robotics).

Imaging was performed using an automated Olympus SpinSR system using a Hamamatsu flash Orca 4.0 camera and a CSUW1-T2SSR SD Yokogawa spinning disk Unit with a 50μm pinhole disk. Images were acquired using a 60× air lens NA 0.9 (Olympus), 100mW 488nm OBIS LX laser system (Coherent), GFP Filter set [EX470/40, EM525/50] (Chroma).

Images were manually inspected using Fiji-ImageJ software (Schindelin et al., 2012).

### High-throughput proximity biotin ligation assay

#### Culturing, induction, and protein extraction

Cells were moved from agar plates into liquid 384-well plates using a RoToR bench-top colony arrayer (Singer Instruments). Liquid cultures were grown overnight in low biotin 0.512nM d-biotin (sigma) SD medium in a shaking incubator (LiCONiC Instruments) at 30°C. A Tecan freedom EVO liquid handler (Tecan) which is connected to the incubator, was used to back-dilute the strains to ∼0.25 OD_600_ in plates containing no biotin SD media supplemented with 1mM Auxin (sigma). Plates were then transferred back to the incubator and were allowed to grow for 4hour at 30°C to reach logarithmic growth phase The liquid handler was then used to prepare samples to measure OD using a plate reader (Tecan) and to add high biotin media 30nM d-biotin(sigma) to the samples which were then incubated at RT for 60min. Then the plates were centrifuged at 3000g for 3min using a robotic centrifuge (Hettich), the media were removed and the cells were resuspended with 25µl of lysis buffer containing 0.1M NaOH, 0.05M EDTA, 2% SDS, 100mM DTT, 1mM PMSF(sigma), orange G dye (adopted form (von der Haar, 2007)) and moved to 384 wells PCR plate (Eppendorf) with reusable lid (4tetude), the PCR plates were moved to a robotic thermal cycler (Inheco) and the samples were incubated at 90° for 10min, neutralized using 5µl of 0.5M acetic acid and incubated again at 90° for 10min

#### Blotting, development, and image acquisition

The lysate was blotted to a nitrocellulose membrane (pall) using the RoToR into a 1536 dot array using the 384 long pads (Singer Instruments) calibrated to move 200pL of liquid. The membrane was dried overnight at RT and blocked for 30min with 2%(W/V) BSA in PBS (Sigma), incubated while shaking ON at 4°C with rabbit anti-Histone H3(Abcam) antibody used as a control, washed, developed using goat anti-rabbit -IRDye680RD antibody (ab216777; Abcam), and streptavidin-Alexa Fluor790(S11378; Invitrogen), on an odyssey LI-COR imaging system.

### Computational analysis

#### Dot-blot analysis

Image analysis was performed using a version of Fiji-ImageJ (Schindelin et al., 2012) plugin Protein Array Analyzer (G. Carpentier) modified to retrieve the Region Of Interest (ROI) for each dot in the array. A set of MATLAB scripts (available at https://github.com/Maya-Schuldiner-lab/Cel-lctiv) were used to measure the signal for each channel for each dot, normalize the signal using the loading control and the three biological repeats and compare the normalized signal for each set of lysates. A set was excluded in several cases: the repeats showed a standard deviation higher than 3, the normalized signal was with a Z score of 10 standard deviations lower than the rest of the membrane, or one of the strains were missing to begin with as indicated by the pre-lysis OD measurement. A protein was designated as having an interaction preference if a signal with a fold change greater than two was found.

#### SP hydrophobicity analysis

Hydrophobicity analysis was performed using the Kyte-Doolittle hydrophobicity scale MATLAB function (Kyte & Doolittle, 1982).

#### GO term analysis

GO term analysis was performed using Yeastmine (Cherry et al., 2012; Engel et al., 2014) GO term enrichment analysis with Holm-Bonferroni correction. For the microscopy screen, the background set was defined as all genes included in the SWAT collection (Weill et al., 2018; Yofe et al., 2016), and for the interaction screen, the background was defined as all genes included in the AviTag library (Fenech et al., 2022) and that were not excluded as part of the analysis.

## Acknowledgements

We thank Dr. Einat Zalckvar, Dr. Yury Bykov, Dr. Emma Fenech, Dr. Sarah Haßdenteufel and Rosario Valenti from the Schuldiner lab for critical feedback of this manuscript. We are grateful to Dr. Yoav Peleg and Prof. Itay Onn for plasmids. The project was supported by the European Research Council Consolidator Grant OnTarget 864068, an Israel Science Foundation grant (ISF 760/17) and the generous support of the Kekst Family Institute for Medical Genetics. The robotic system of the Schuldiner lab was purchased through the kind support of the Blythe Brenden-Mann Foundation. Maya Schuldiner is an incumbent of the Dr. Gilbert Omenn and Martha Darling Professorial Chair in Molecular Genetics.

## Author contributions

N.C designed, performed and analysed experiments. N.A designed and conceptualized experiments. N.C and M.S wrote the manuscript which all authors read and provided feedback on. M. Schuldiner supervised the work and secured funding.

